# Post-translational modifications via serine/threonine phosphorylation and GpsB in *Streptococcus mutans*

**DOI:** 10.1101/2025.07.25.666849

**Authors:** Sangam Chudal, Courtney Dover, Tiffany Haydt, Shawn M. King, Robert C. Shields

## Abstract

Post-translational modifications (PTMs), such as protein phosphorylation, are critical regulators of bacterial physiology. Here, we present the first comprehensive phosphoproteomic analysis of *Streptococcus mutans*, revealing extensive *O*-phosphorylation under non-stressed conditions. Using tandem mass tag (TMT)-based mass spectrometry and phosphopeptide enrichment, we identified 231 high-confidence phosphosites on 131 proteins, representing approximately 6.7% of the detected proteome. These phosphorylated proteins were enriched in pathways related to translation, carbohydrate metabolism, and the cell cycle, suggesting a broad role for *O*-phosphorylation in core cellular functions. To define the functional roles of the sole serine/threonine protein kinase (PknB) and phosphatase (PppL) encoded by *S. mutans*, we analyzed phosphoproteomic and proteomic changes in Δ*pknB* and Δ*pppL* mutants. These mutants exhibited widespread alterations in protein abundance and phosphorylation, revealing overlapping but distinct sets of putative kinase and phosphatase substrates, including DivIVA, MapZ, MltG, and ribosomal proteins. Notably, we discovered that repression of *gpsB*, a predicted PknB binding partner, causes lethal defects that can be rescued by a suppressor mutation (G98R) in *pppL*. This mutation restores phosphorylation of DivIVA, suggesting that GpsB regulates the PknB/PppL signaling axis to maintain appropriate phosphorylation of essential targets. This work highlights conserved and unique features of bacterial phospho-signaling and provides a foundation for future studies on PTM-mediated regulation in *S. mutans*.

## Introduction

Over the past two decades, there has been a growing understanding of the vital role that post-translational modifications (PTMs) have for bacterial physiology (comprehensively reviewed by Macek et al., (1)). This growing body of knowledge highlights that bacteria, despite their simpler cellular organization compared to eukaryotes, employ a sophisticated array of PTMs to fine-tune their cellular processes and adapt to dynamic environmental cues. A critical aspect of PTMs is that these modifications are generally not present on all molecules of a given protein. This differential modification adds another level of regulation and ability to respond to external stressors (2). For instance, a protein might exist in both a modified and unmodified state, with each state exhibiting distinct activities or cellular localizations. This heterogeneity allows bacterial populations to rapidly and efficiently adjust their proteome in response to nutrient availability, osmotic stress, pH changes, temperature fluctuations, the presence of antibiotics, or host immune responses (3). Understanding bacterial PTMs is crucial not only for deciphering fundamental bacterial biology but also for identifying novel targets for antimicrobial drug development.

Protein phosphorylation is a major source of PTMs in bacteria (reviewed by Frando and Grundner (4)). In bacteria, these modifications have been detected at histidine, aspartate, arginine, cysteine, serine, threonine, and tyrosine residues (5–8). The best characterized, and probably most common, type of phosphorylation in bacteria occurs via two component systems (TCS) where a signal is transmitted via histidine autophosphorylation, and this leads to changes in gene expression (9). Despite their widespread importance, the transient and inherently unstable nature of histidine phosphorylation within TCS makes their *in vivo* detection a considerable challenge (10). In contrast, *O*-phosphorylation, where a phosphate group is added to the hydroxyl group (-OH) of serine, threonine, or tyrosine residues in a protein is more stable and can be detected via mass spectrometry approaches. However, despite technological advancements, comprehensive global phosphoproteome studies specifically focused on elucidating the full extent and significance of *O*-phosphorylation in bacteria remain relatively few. This highlights a significant knowledge gap and underscores the need for more extensive investigations to fully appreciate the crucial role of *O*-phosphorylation in bacterial physiology, virulence, and adaptation.

In bacteria, *O*-phosphorylation is carried out by eukaryotic-like serine/threonine protein kinases (STKPs) and bacterial tyrosine-kinases (BYKs). Although widespread, the total number and distribution of these proteins is highly variable. For example, *Mycobacterium tuberculosis* encodes eleven STKPs (11), *Bacillus subtilis* encodes four STKPs (12), and most streptococci only encode a single STKP (13–15). In the dental caries pathogen, *Streptococcus mutans*, there is a single STKP known as PknB and it lacks a BYK. Mutagenesis of *pknB* and/or the serine-threonine protein phosphatase (STP) known as *pppL* has widespread phenotypic effects in *S. mutans* (16–18). Other than these phenotypic studies, a comprehensive phosphoproteome of *S. mutans* has not been characterized. Here, we uncovered the *O*-phosphorylation proteome of *S. mutans* using mass spectrometry (MS). To gain a deeper understanding, we also investigated the phosphoproteomes of *S. mutans* mutants deficient in *pknB* and *pppL* to identify putative cellular substrates of these proteins. Lastly, we characterize the impact that interfering with a possible PknB binding partner, GpsB, has on *O-*phosphorylation. These data uncover the critical role that *O*-phosphorylation has in *S. mutans*, laying a foundation from which to understand many of the predicted phosphorylated proteins and protein-protein interactions.

## Results

### Phosphoproteomics reveals the *S. mutans O*-phosphoproteome in non-stressed conditions

To comprehensively detect phosphorylated serine, threonine, or tyrosine residues we employed a multiplexed tandem mass tags (TMT) MS identification and quantification methodology. Our workflow began with collecting cell pellets from *S. mutans* UA159 cultured to an OD_600_ ∼0.4-0.6. Bacteria generally have fewer phosphorylated proteins than eukaryotes, so we collected cell pellets from 15 mL of culture to increase the likelihood of capturing low abundance phospho-proteins. In total, our analysis was carried out across fifteen independent replicates. For each replicate, the cells were lysed and proteins extracted after which the proteins were digested. After digestion, peptides from each sample were individually labelled with TMT tags and phosphorylated proteins were enriched using immobilized metal ion affinity chromatography (IMAC) resins. For these highly complex samples, the peptide mixtures were then fractionated by reverse-phase chromatography to reduce sample complexity and improve the depth of phosphoproteome coverage. Samples were then loaded onto an autosampler, concentrated with a trapping column, and then separated by an acetonitrile gradient on an analytical column. Separated peptides were then analyzed by MS, MS/MS, and the data was analyzed using MaxQuant software.

Across the fifteen replicates, MS measurements detected 1572 proteins, corresponding to 80% of the *S. mutans* UA159 protein coding genes. From these fifteen samples we were able to detect 231 pSer/pThr/pTyr phosphosites from 131 unique proteins. To strictly assign phosphorylation status, we only considered these sites to be phosphorylated if they met a criterion of a localization probability (LP) score ≥ 0.75. The median LP score for these phosphosites was 0.999 and the mean LP score was 0.973, indicating a high confidence of phosphorylation for virtually all the phosphopeptides. Taken together, *O*-phosphorylation corresponds to 6.7% of the total number of proteins that were detected across the fifteen samples. Within this set of peptides, 49.4% were phosphorylated on a threonine, 42.4% on a serine, and 8.2% on a tyrosine residue (**Figure 1A**).

**Figure 1:**
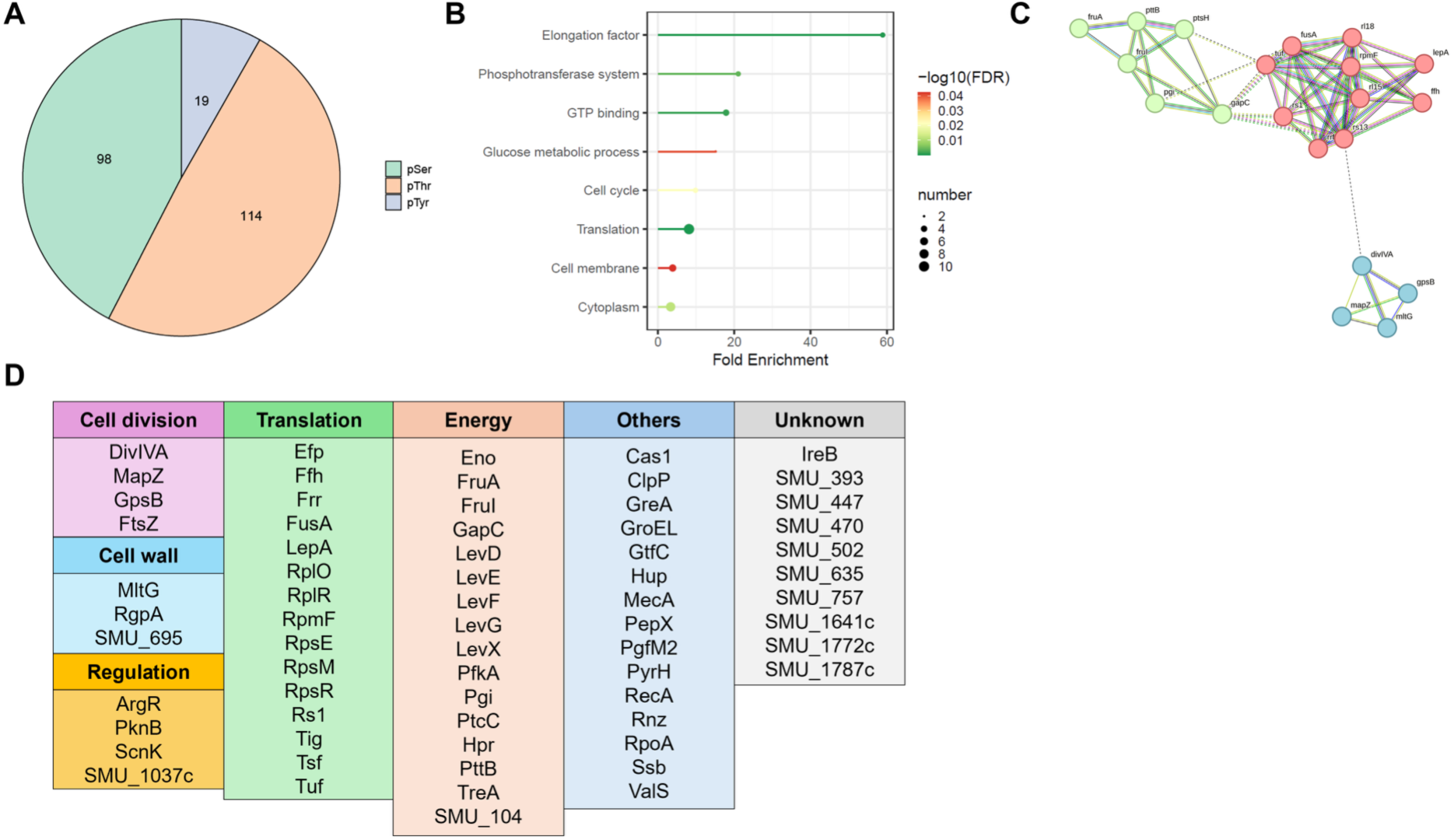
The *S. mutans* phosphoproteome. (**A**) Relative share of phosphorylated serine, threonine, and tyrosine detected in *S. mutans*. (**B**) Lollipop plot highlighting that *O*-phosphorylated proteins have an enrichment for carbohydrate metabolism, translation, and the cell cycle. (**C**) STRING protein-protein interactions between predicted phosphorylated proteins. The lines connecting proteins represent functional associations that are either experimental or text-mining based. There are three distinct protein hubs related to carbohydrate metabolism, translation, and the cell cycle. (**D**) Summary of the proteins with detected phosphorylated serine, threonine, and tyrosine amino acid residues.

A list of all the detected phosphopeptides is available in **Table S1**, and this table also describes the number of samples (out of fifteen) that a phosphopeptide was detected in. Notably, only 23 phosphopeptides were detected in 100% of the samples, and 51 were detected in greater than 50% of the samples. These likely represent proteins/pathways for which *O*-phosphorylation is critical for function in the conditions tested, and thus detection is stable. We performed protein/gene-set enrichment for these phosphorylated proteins to determine if particular biological processes are enriched for these proteins. **Figure 1B** shows that there is enrichment for proteins with functions related to translation, carbohydrate metabolism, the cell cycle, and that these proteins are enriched both within the cytoplasm and the cell membrane. Known and putative interactions between these phosphorylated proteins is shown in **Figure 1C**, with distinct groupings for translation, carbohydrate utilization, and cell cycle proteins.

Several components of the protein translation apparatus appear to be phosphorylated. We detected phosphorylation of Ffh, a component of the bacterial signal recognition particle (SRP), at Ser397 (LP = 0.997). Ribosomal proteins RpmF at Thr32 (LP = 0.958), RpsM at Ser50 (LP = 0.834), and Rs1 at Thr198 (LP = 0.991) are phosphorylated. Elongation factor Tu (also known as EF-Tu and Tuf) is phosphorylated at Ser52 (LP = 0.982). Components of the cell cycle that are phosphorylated include MapZ which was detected at Thr18 (LP = 0.999), and Thr127 (LP = 0.983). The peptidoglycan hydrolase, MltG, which has a role in determining glycan chain lengths is phosphorylated at Thr102 (LP = 0.928), Thr166 (LP = 1), and Thr191 (LP = 1). Several carbohydrate utilization proteins also appear to be phosphorylated, including the histidine-containing phosphocarrier protein (HPr; also known as PtsH), at Thr12 (LP = 1), which has a key role in the phosphoenolpyruvate-dependent sugar phosphotransferase system (PTS).

### Identification of the serine/threonine kinase and phosphatase cellular substrate network

Next, we sought to characterize the impact on the proteome and the phosphoproteome of decreased and increased levels of *O*-phosphorylation. Based on prior characterization (16) and genome mining, *S. mutans* UA159 has a single STKP (known as PknB) and a single phosphatase (known as PppL). We anticipated that deletion of *pknB* would lead to a detectable decrease in *O*-phosphorylation and that deletion of *pppL* would have the opposite effect. Mutant strains of *pknB* and *pppL* were previously generated as part of a study exploring genetic competence regulation in *S. mutans* (19). Five replicates of either Δ*pknB* or Δ*pppL* were cultured to mid-log, along with five replicates of *S. mutans* UA159 for each comparison. Proteins and phosphoproteins were then detected as described in the Materials and Methods.

We began by investigating changes in the proteome for both *S. mutans* Δ*pknB* and Δ*pppL* compared to *S. mutans* UA159. When comparing between wild-type and mutant strains, we considered protein abundance changes with a p-value < 0.05 and an increase or decrease equal to or greater than a log2 fold-change of 1. In total, 223 and 485 proteins met this criteria for Δ*pknB* and Δ*pppL* respectively (**Figure 2**; **Tables S2 and S3**). The broad changes in the proteome for both the *pknB* and *pppL* mutants is consistent with the genetic competence, biofilm, cell wall, and transcriptional phenotypes that have been described for these strains (16). Given that 10-25% of the proteome was altered in these strains, according to our criteria, we sought to quantify the pathways that are most enriched to potentially highlight critical biological processes that are disturbed. Both up- and down-regulated proteins in Δ*pknB* and Δ*pppL* mutants were significantly enriched for genes from specific biological pathways (**Figure 2**). Up-regulated proteins in the Δ*pknB* mutant were enriched for the TnSmu1 mobile genetic element, nucleotide metabolism, and a pathway related to ATP binding cassette (ABC) transporters. Down-regulated proteins in the Δ*pknB* mutant were enriched for secreted/cell-well-bound proteins, carbohydrate transport, lactose metabolism, aromatic amino acid biosynthesis, lipoteichoic acid metabolism, and glycosyltransferases (Gtfs, including GtfB, GtfC, and GtfD). For the Δ*pppL*mutant, we observed up-regulated proteins in pathways related to organic anion transport, AI-2E family transporters, and proteins with a transmembrane helix. Down-regulated proteins in the Δ*pppL* mutant were enriched for peptidoglycan synthesis, chorismate metabolism, DNA replication, DNA repair, aromatic amino acid biosynthesis, amino acid biosynthesis, and ribosomal proteins/regulation of translation.

**Figure 2:**
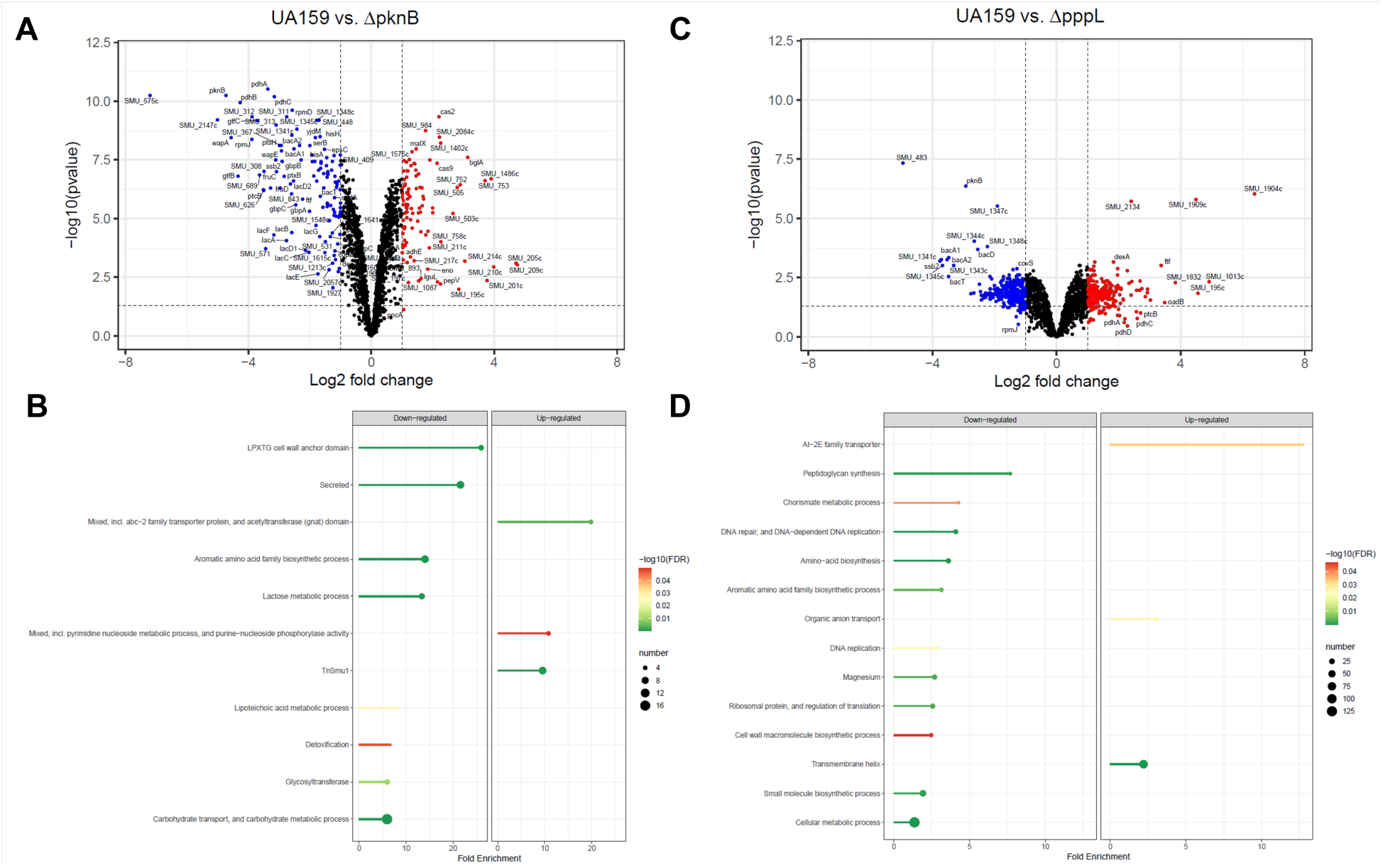
Proteomics analysis of *S. mutans* Δ*pknB* and Δ*pppL*. (**A**) Volcano plot comparing the proteome of *S. mutans* UA159 with Δ*pknB*. The horizontal and vertical hatched lines show significance cut-offs for the data (p-value < 0.05 and log2 fold change greater or less than 1). (**B**) Lollipop plot highlighting biological pathways that were significantly enriched or depleted in Δ*pknB* compared to *S. mutans* UA159. (**C**) Volcano plot comparing the proteome of *S. mutans* UA159 with Δ*pppL*. The horizontal and vertical hatched lines show significance cut-offs for the data (p-value < 0.05; log2 fold-change >1 or <-1). (**D**) Lollipop plot highlighting biological pathways that were significantly enriched or depleted in Δ*pppL*compared to *S. mutans* UA159.

Next, we analyzed the phosphoproteome of the Δ*pknB* and Δ*pppL* mutants compared to *S. mutans* UA159. In total, we detected 42 and 234 phosphosites in our analysis of Δ*pknB* and Δ*pppL* mutants respectively (**Tables S4 and S5**). In Δ*pknB* a total of 22 phosphosites showed a significantly altered abundance compared to *S. mutans* UA159 (p-value < 0.05; log2 fold change > 1). Of these, 14 had decreased abundance (including phosphosites only detected in UA159 and not in Δ*pknB*) and 8 had increased abundance compared to wild-type samples (including phosphosites only detected in Δ*pknB* and not in UA159). For Δ*pppL* we observed a total of 126 phosphosites with significantly altered abundance of which 36 had decreased abundance and 90 had increased abundance (including phosphosites only detected in Δ*pppL* and not in UA159). These broad observations are consistent with the hypo- and hyperphosphorylation that would be expected in Δ*pknB* and Δ*pppL* mutants respectively.

Next, we analyzed proteins with a reduced phosphorylation level in the Δ*pknB* strain and increased phosphorylation level in the Δ*pppL* strain. As shown in **Figure 3**, there are twenty six proteins where phosphosites were detected in both the Δ*pknB* and Δ*pppL* mutant strains. The expected hyperphosphorylation was apparent for Δ*pppL* as several additional phosphosites were detected in these proteins compared to Δ*pknB*. For example, seven phosphosites were detected in DivIVA in Δ*pppL* samples, and only two in the Δ*pknB samples.* To determine putative PknB cellular substrates we scrutinized for phosphosites detected in all wild-type samples that were absent or significantly reduced (log2 fold change < −5) in the Δ*pknB* mutant strain. Our conservative list of probable PknB phosphorylation targets is six proteins and a total of eleven phosphosites: DivIVA (Thr232 and Thr262), MapZ (Thr18 and Thr127), RpmF (Thr32), MltG (Thr102, Thr166, and Thr191), an IreB-like protein (SMU_2079c; Thr4 and Thr7), and PgfM2 (Thr16). Although direct evidence is required for conclusive identification of these PknB targets there is an overlap with predicted *S. pneumoniae* StkP substrates for DivIVA, MapZ, MltG, and Spr0175 (IreB-like protein). Likely *S. mutans* unique PknB substrates include RpmF, the L32 ribosomal protein, and PgfM2, a transporter that is part of the Pgf glycosylation machinery (20). We consider our list of putative PknB targets to be conservative because we used non-stressed growth conditions. In the event of stressful conditions, external signals gathered by PknB probably alter phosphorylation levels of specific proteins. Nevertheless, strong evidence suggests that *S. mutans* PknB likely has a direct role in modifying the activity of proteins involved in cell division, peptidoglycan synthesis, translation, and glycosylation (**Figure 4**).

**Figure 3:**
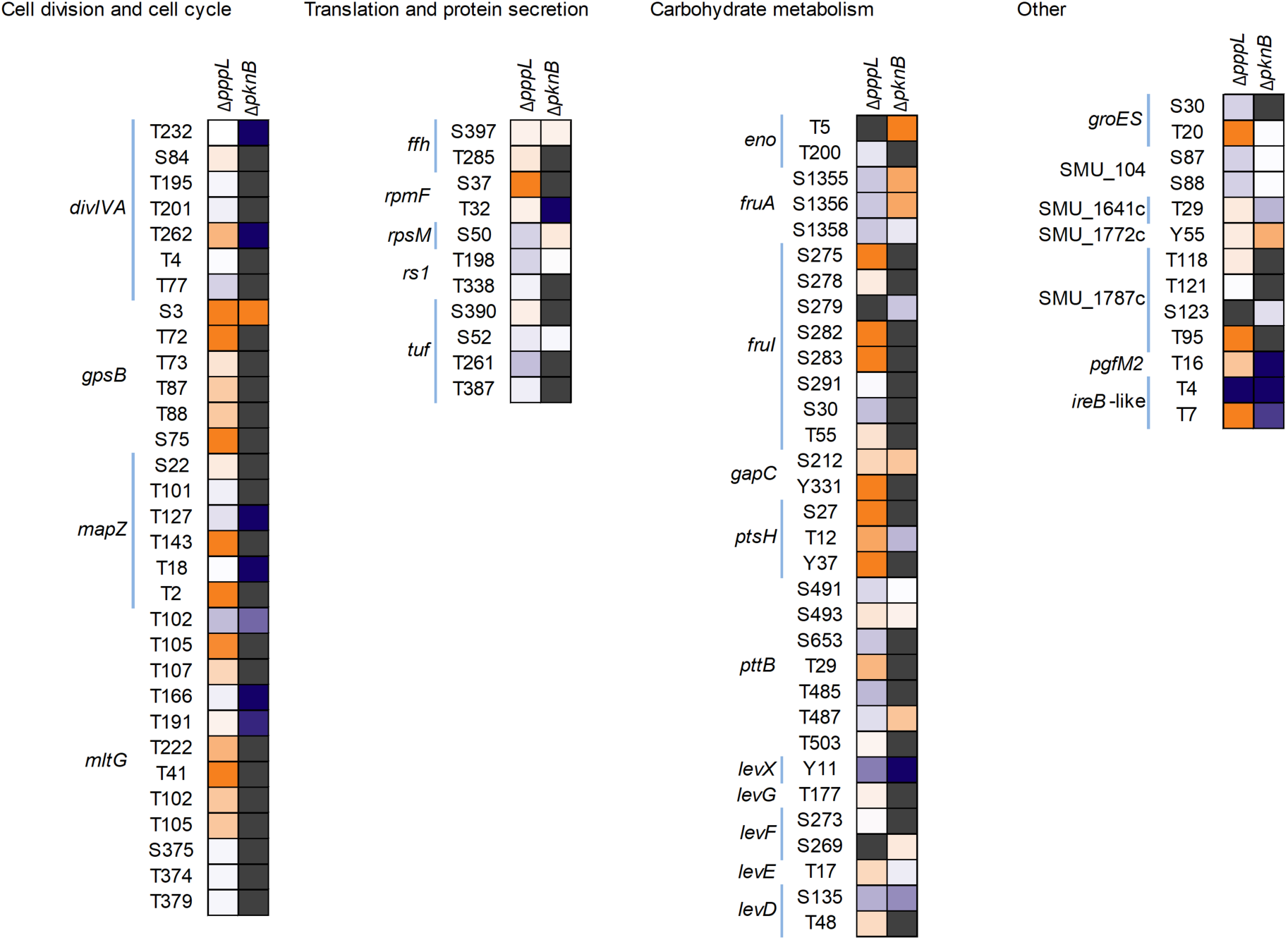
Heatmap visualizing hyper- and hypophosphorylation of serine, threonine, and tyrosine residues in Δ*pknB* and Δ*pppL* mutant strains. Phosphorylated proteins are sorted according to their biological function. Orange indicates increased phosphorylation compared to the control strain, and blue indicates reduced phosphorylation compared to the control strain. Grey indicates that phosphorylation was not quantified at that residue for that strain.

**Figure 4:**
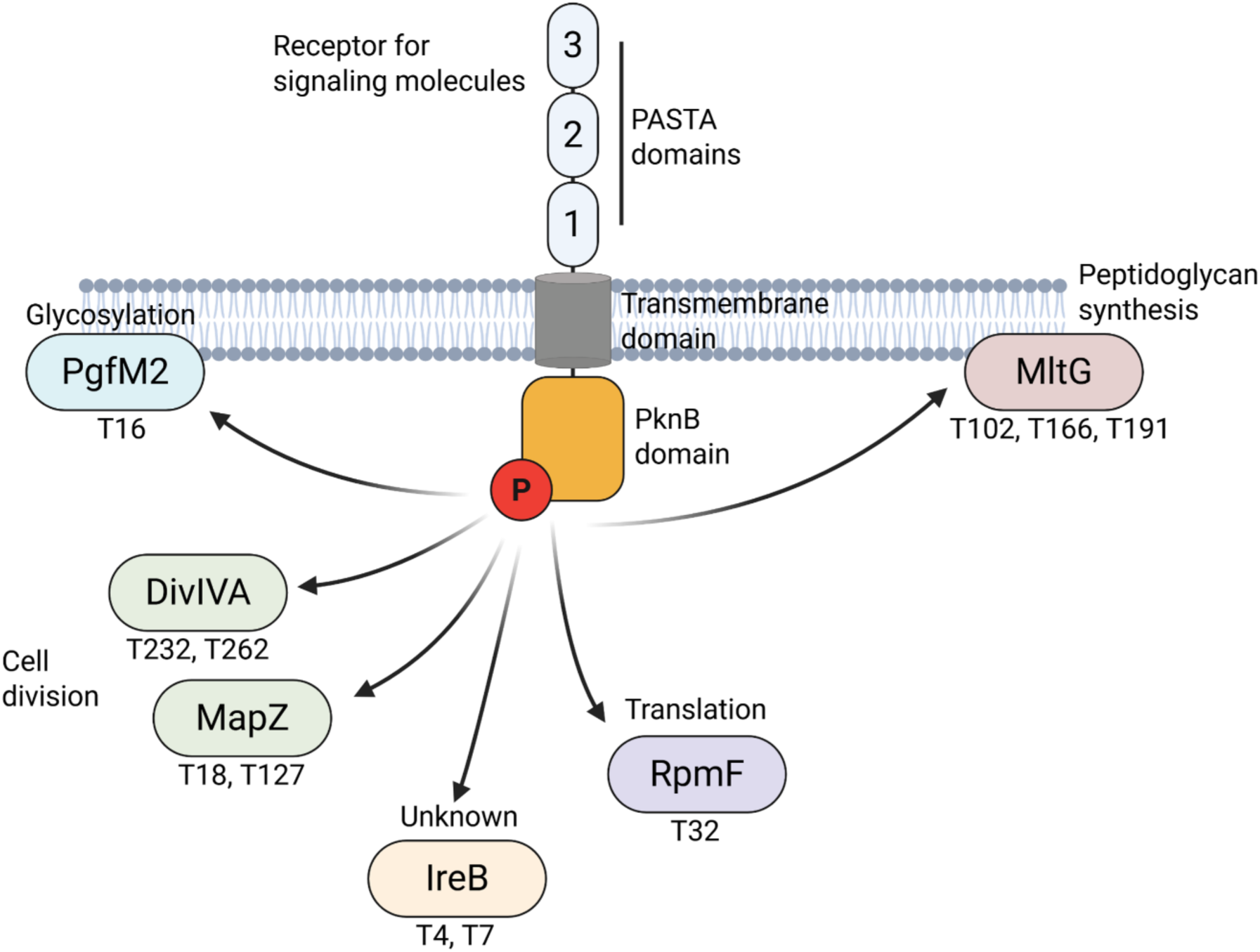
Diagram of putative *S. mutans* PknB cellular substrates. PknB PASTA domains stimulate PknB autophosphorylation in response to an external signal (likely muropeptides). Putative targets of PknB, as determined through phosphoproteomics, are shown with specific amino acid residues highlighted. PknB phosphorylates proteins involved in cell division, peptidoglycan synthesis, translation, and glycosylation.

While putative STKP targets have been detected and characterized for several Gram-positive microorganisms, there has been considerably less attention on possible phosphatase targets. With that in mind, we wanted to identify putative PppL targets, which we defined as pSer/pThr/pTyr residues detected in at least three out of five Δ*pppL* samples with no detection in wild-type samples (LP score ≥ 0.75). Notably, a total of fifty-one phosphosites within forty-three proteins met these criteria (**Table S6**). These phosphosites/proteins can be split into two broad groups: (1) proteins with increased phosphosites in Δ*pppL*, and (2) proteins with phosphosites only in Δ*pppL*. Example proteins for the first group include MapZ (Thr2, Thr143), FruI (Ser275, Ser282, Ser283), GapC (Tyr331), GpsB (Ser3, Ser75), GreA (Ser79), GroEL (Thr430), Hup (Thr18), and HPr/PtsH (Tyr37). The following phosphosites in DnaK (Thr430, Tyr104), Glk (Ser210, Ser283), ProS (Ser286), QueF (Thr10), RplX (Ser66), RpmA (Thr41), RpsJ (Ser35), and SecY (Tyr347), were only detected in the Δ*pppL* mutant strain. There are several proteins that are predicted to be dephosphorylated by PppL that were not predicted to be phosphorylated by PknB (e.g. GroES [Thr20] and SecY [Tyr347]). However, it is conceivable that our predicted PknB cellular substrates is a minimal set, given that cells were grown in non-stressed conditions. PknB activity, and the number of cellular substrates, may increase depending on the environmental conditions that a cell encounters (deletion of *pppL* itself is a stress for the cell). Hyperphosphorylation sites in a Δ*pppL* strain may also indicate *O*-phosphorylation sites that are less stable in wild-type cells because they are rapidly dephosphorylated by PppL. Alternatively, loss of *pppL* could cause PknB to become hyperactive/more promiscuous and these phosphosites are not biologically relevant. Despite the apparently differential phosphorylation patterns comparing Δ*pknB* or Δ*pppL* the results are consistent with PppL participating in biological pathways that overlap considerably with PknB (carbohydrate metabolism, translation, and the cell cycle).

### Functional consequences of repressing GpsB, a predicted PknB partner

The <u>g</u>uiding <u>P</u>BP1 and <u>s</u>huttling (GpsB) protein has been shown to have an important role in cell division for Gram-positive microorganisms (21). Of particular relevance to this study, is findings that GpsB is phosphorylated by STKPs (22, 23) and it is required for STKP-mediated phosphorylation of DivIVA, MapZ, and other proteins (24). According to our phosphoprotemics analysis, *S. mutans* GpsB is putatively phosphorylated at Thr72, Thr73, Thr87, and Thr88 in wild-type cells. We also observed phosphorylation at Ser3 in both Δ*pknB* and Δ*pppL* mutant samples and Ser75 in Δ*pppL* mutant samples. GpsB phosphosites in other microorganisms include Thr88 (*Listeria monocytogenes*) (25), Thr79/Thr83 (*Streptococcus pneumoniae*) (23, 26), Thr75 (*Bacillus subtilis*) (22), Ser80/Thr84 (*Enterococcus faecalis*) (27). Alignment of *S. mutans* GpsB with these GpsB proteins is shown in **Figure S1** with the detected phosphosites highlighted.

Our initial goal here was to characterize the impact that loss of *gpsB* has on *S. mutans* physiology. We previously determined that *gpsB* is an essential gene (cannot be deleted), and that repression of *gpsB* with a Clustered Regularly Interspaced Short Palindromic Repeats interference (CRISPRi) tool causes cell division defects (28). Using the same CRISPRi strain (CRISPRi*^gpsB^*), we explored *gpsB* phenotypes in more detail. First, we serially plated the CRISPRi*^gpsB^*strain on eleven different conditions to mimic stress across multiple cellular pathways (Figure 5A). In these experiments, 0.5% xylose is included, as this leads to induction of dCas9 (dead Cas9) and a concomitant repression of *gpsB*. The CRISPRi*^gpsB^*strain grew poorly across all the conditions but there was visually less growth during acid (BHI pH 5.5) and transcription (rifampicin) stress. Next, we used transmission electron microscopy (TEM) to observe the cell morphology of control and *gpsB* repressed samples (**Figure 5B**). There was a significant increase (p < 0.001) in the mean cell width of *gpsB* repressed (589 nm) versus *gpsB* non-repressed cells (429 nm) (**Figure 5C**). As a result, *gpsB* repressed cells have a rounder appearance compared to control cells.

**Figure 5:**
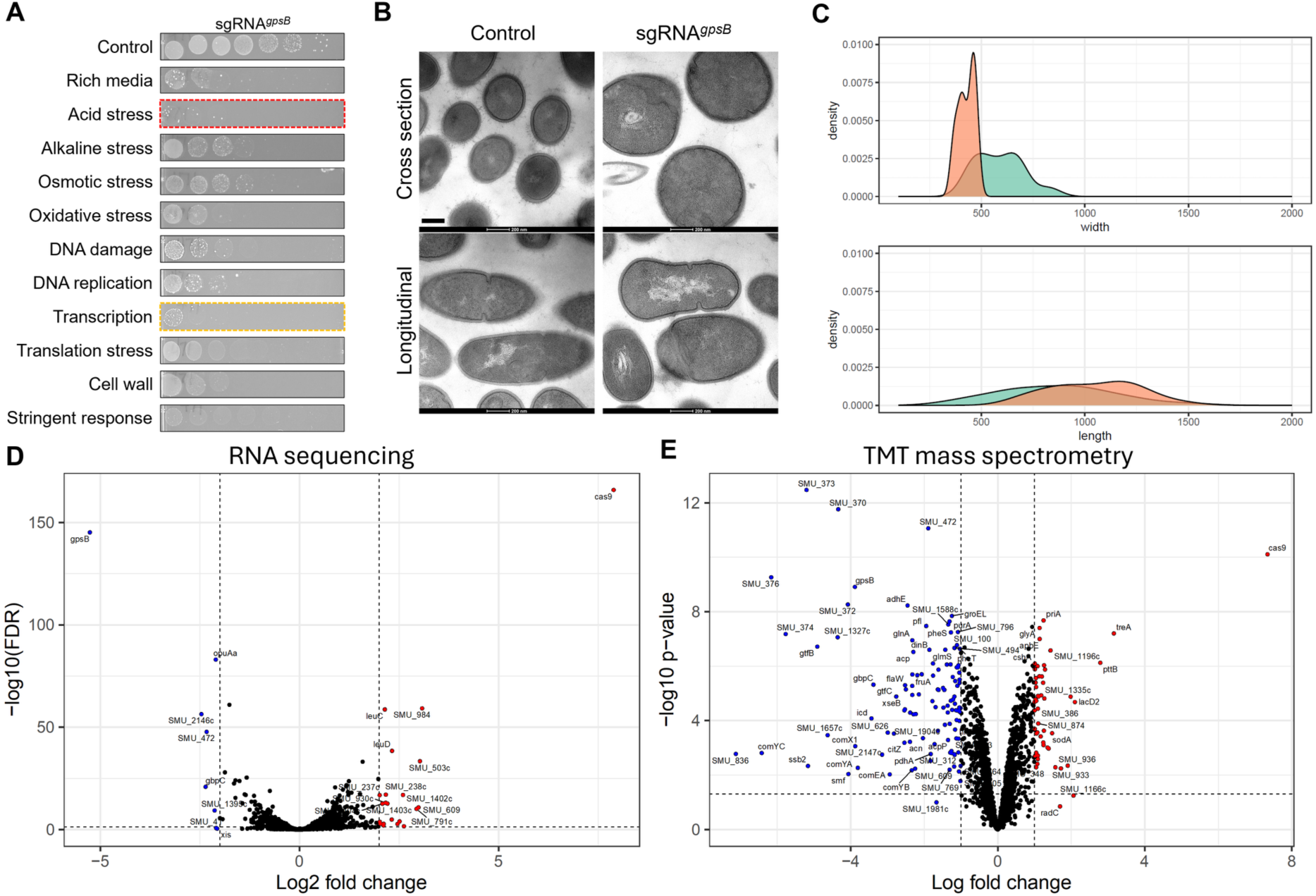
Repression of *S. mutans gpsB* inhibits growth, alters cell morphology, and causes broad changes in the proteome. (**A**) A CRISPRi strain targeting *gpsB* was serially diluted and spot plated onto BHI agar plates under various stress conditions. Acid stress, BHI pH 5.5; alkaline stress, pH 8.5; osmotic stress, 2% NaCl; oxidative stress, 0.5 mM H_2_O_2_; DNA damage, 10 ng/mL mitomycin C; DNA replication, 1 µg/mL ciprofloxacin; transcription stress, 0.075 µg/mL rifampicin; translation stress, 1.5 µg/mL chloramphenicol; cell wall stress, 2 µg/mL ampicillin; stringent response, 0.015 µg/mL mupirocin. Experiments were repeated four times with similar results. (**B**) Ultrastructural analysis of *gpsB* repression using transmission electron microscopy. (**C**) Density plot of ImageJ quantification of cell widths and lengths comparing the control (orange) with CRISPRi*^gpsB^* (green). (**D**) Volcano plot of RNA-seq transcriptome data showing gene expression values for *gpsB* repressed relative to *gpsB* non-repressed strains cultured in chemically defined medium. Differentially expressed genes (FDR ≤0.05; log2 fold-change >2 or <-2) are defined by the hatched lines. (**E**) Volcano plot of whole-cell proteomics showing protein expression values for *gpsB* repressed relative to *gpsB* non-repressed strains cultured in rich media. Differentially expressed proteins (log2 fold-change >1 or <-1; adjusted p-value ≤0.05) are defined by the hatched lines.

To assess the global importance of GpsB we conducted whole-cell proteomics, and transcriptomics. To quantify changes in gene expression, we conducted RNA-seq comparing the transcriptomes of CRISPRi*^gpsB^* without and with (causing *gpsB* repression) xylose. Genes with significant changes in expression were defined as those with a log2 fold change > 2.0 and a false discovery rate (FDR) < 0.05. A total of 8 genes were down-regulated, and 21 genes were up-regulated when gpsB was repressed (**Table S7**). There was evidence that the CRISPRi system was working as expected as *gpsB* expression was down-regulated by 38-fold, and *cas9* was up-regulated 236-fold. Overall, the changes in the *S. mutans* transcriptome in *gpsB*-repressed cells were modest. Next, we used TMT quantitative proteomics to compare proteins between CRISPRi*^gpsB^* without and with xylose. A total of 1,523 proteins were detected from *S. mutans* cell lysates. Proteins with significant changes in abundance were defined as those with a log2 fold change > 1.0 and an adjusted P value < 0.05. Using those criteria, 48 proteins were up-regulated, and 120 proteins were down-regulated with *gpsB* repression (**Table S8**). Down-regulated proteins included several that are secreted/cell wall associated (GtfB, GtfC, WapE, GbpC, DexA, SpaP, FruA, WapA), involved with DNA transformation (Smf, ComYC, ComYB, ComYA, ComEA, ComX), and histidine metabolism (HisF, HisA, HisH, SerB, HisD, HisZ, HisC). For up-regulated proteins there were few enriched pathways, except for poorly characterized transporter systems (SMU_1217c, SMU_932, SMU_933, SMU_936, SMU_1067c, SMU_1068c, SMU_1070c).

### The essentiality of *gpsB* can be reversed via the acquisition of suppressor mutations in the *pppL* phosphatase

Next, we hypothesized that *gpsB* gene replacement in tandem with suppressor mutation acquisition might restore viability for mutant strains, and that this could highlight critical *gpsB* interacting genes/pathways. This would be consistent with prior studies showing that the essentiality of *gpsB* can be bypassed in both *S. pneumoniae* and *L. monocytogenes* through the acquisition of suppressor mutations (29, 30).

To test this, we designed a construct to replace *gpsB* with an antibiotic resistance cassette and transformed *S. mutans* via genetic competence. Despite several attempts at mutagenesis of *gpsB* we were only ever able to isolate one viable *gpsB* mutant (named Δ*gpsB*_sup_). Replacement of *gpsB* with a kanamycin resistance gene was confirmed by PCR and agarose gel electrophoresis for this single viable mutant. The Δ*gpsB*_sup_ mutant strain grew similar to wild-type *S. mutans* UA159 in rich media (**Figure 6A**). We sequenced the genome of the Δ*gpsB*_sup_ strain and it contained two suppressor mutations. One of the suppressor mutations was within a gene of unknown function, SMU_1232c, and is a single amino acid change (S42R). The other suppressor mutation was also a single amino acid change (G98R) in *pppL* (**Figure 6B**), which is, as described above, a Ser/Thr phosphatase.

**Figure 6:**
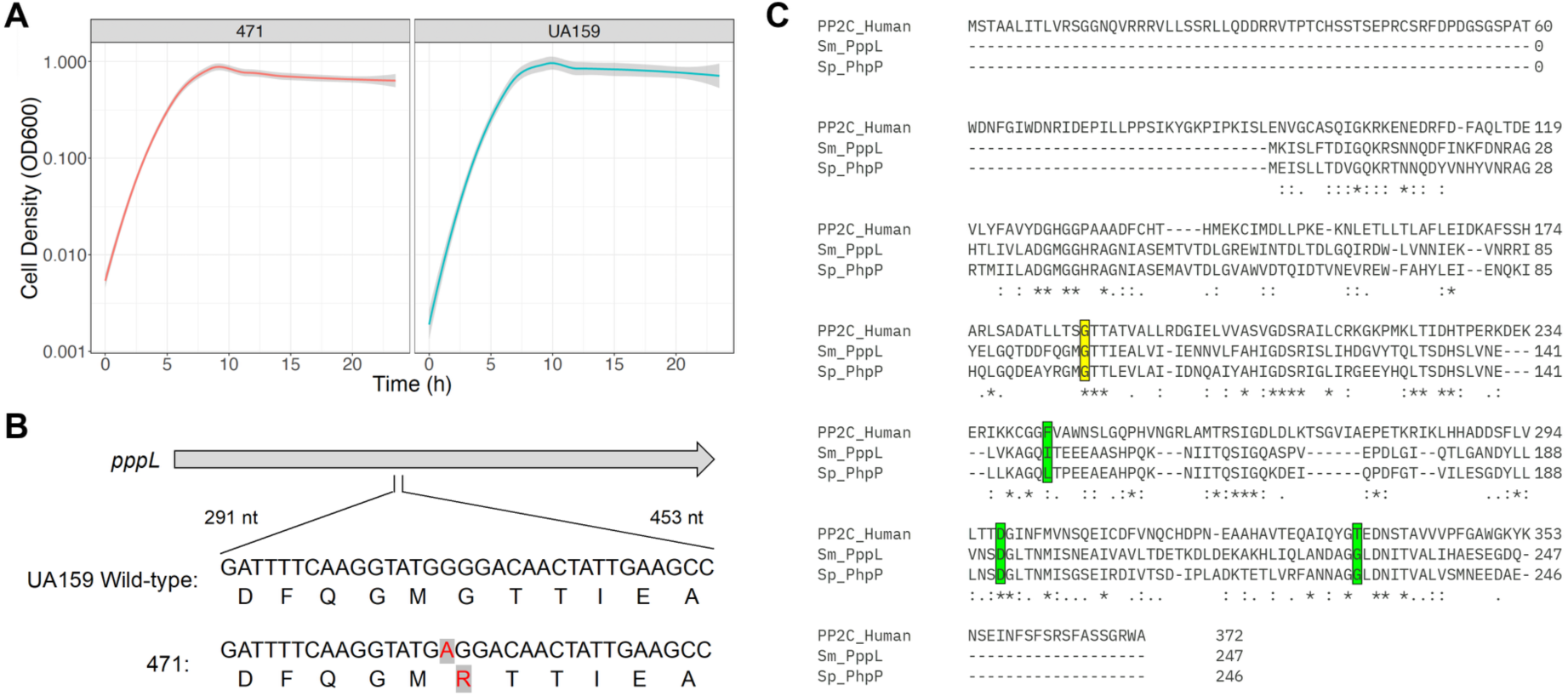
A *pppL* single nucleotide polymorphism suppresses the essentiality of *S. mutans gpsB*. (**A**) Growth analysis of Δ*gpsB*_sup_ and UA159 in a rich medium. Cell density was measured using a Bioscreen C° Pro automated growth system. (**B**) Diagram highlighting the guanine (G) to adenine (A) mutation found in *pppL* after deletion of *gpsB*. This mutation converts glycine to an arginine amino acid residue. (**C**) Alignment of *S. mutans* PppL, *S. pneumoniae* PhpP, and human PP2C serine/threonine protein phosphatase enzymes. Highlighted in yellow is the conserved glycine amino acid residue that is mutated in the Δ*gpsB*_sup_ strain (G98R). Green highlighted regions are those for which *phpP* mutations were found to restore the viability of *gpsB* deletion in *S. pneumoniae* (L148S, D192A, G229D).

Although we cannot rule out that the SMU_1232c has an important role in allowing *gpsB* deletion, of particular interest to us was the mutation in *pppL*. Notably, *gpsB* essentiality can be suppressed by mutations that inactivate the Ser/Thr phosphatase, known as PhpP, in *S. pneumoniae* (30). In *S. pneumoniae* GpsB is required for protein phosphorylation, and mutations that inactivate PhpP restore protein phosphorylation levels for Δ*gpsB* mutants (30). There were several suppressor mutations (D192A, G229D, and L148S) that inactivated PhpP in *S. pneumoniae,* but these are in different locations to the G98R mutation that we observed in the Δ*gpsB*_sup_ strain (Figure 6C). *S. mutans* PppL belongs to a large family of serine/threonine phosphatases, known as the PP2C family (31), that is widespread in bacteria. There are also PP2C phosphatases present in eukaryotic organisms including humans. To determine if the Gly98 residue has a conserved and potentially important role in PppL function, we aligned *S. mutans* PppL, *S. pneumoniae* PhpP, and a human PP2C protein. Our rationale for exploring this conservation centres on the observation that residues that remain constant across species have been maintained because they are important for a protein’s shape and function. As highlighted in Figure 6C, the glycine residue is conserved as part of a triad of amino acids (glycine, threonine, threonine; GTT) across the three species. Although functional studies are required to determine any role of this triad on protein function, the conservation suggests that these residues are important for PppL activity (and related PP2C phosphatases).

### GpsB modifies serine/threonine kinase phospho-signaling

Based on the observation that *gpsB* can be deleted via acquisition of *pppL* mutations, we next wanted to examine *O*-phosphorylation in *gpsB* mutant strains. Our hypothesis, consistent with prior studies in *S. pneumoniae*, was that *gpsB* is essential because it has a critical role in protein phosphorylation and that *pppL* suppressor mutations allow *gpsB* deletion because of hyperphosphorylation. Initially we observed the phosphorylation status of *S. mutans* strains using a phospho-threonine/tyrosine antibody that detects phosphorylated proteins at threonine/tyrosine residues with some cross reaction at phospho-serine residues. However, as shown in **Figure S2**, results with this antibody were mixed. Although there was the expected result of hyper- and hypophosphorylation in Δ*pppL* and Δ*pknB* respectively, relatively few phosphorylated proteins were detected in *S. mutans* UA159. Repression of *gpsB* with CRISPRi gave mixed results, with increased and decreased abundance of phosphorylated proteins compared to control strains. Markedly, the Δ*gpsB*_sup_ had an overall pattern of hyperphosphorylation which would suggest that the G98R *pppL* mutation inactivates phosphatase activity as hypothesized.

To gain a more quantitative and nuanced view of *gpsB* mutant strain phosphoproteomes, we next conducted phospho-TMT analysis of proteins collected from CRISPRi*^gpsB^* (with and without xylose) and Δ*gpsB*_sup_ (**Tables S9 and S10**). A heatmap diagram depicting increased and decreased abundance of phosphorylation of proteins in both strains is shown in **Figure 7**. Focusing on the CRISPRi*^gpsB^* results, we observed that several predicted PknB cellular substrates (**Figure 4**) had significantly decreased phosphorylation (p-value < 0.05; log2 fold-change < 1; or absent in four out of five samples) at specific amino acid residues: DivIVA (Thr232), MapZ (Thr127), RpmF (Thr32), MltG (Thr191), and PgfM2 (Thr16). The only predicted PknB substrate that we did not observe a significant decrease in phosphorylation was for SMU_2079c (an IreB-like protein). By contrast, increased phosphorylation (p-value < 0.05; log2 fold-change > 1) was observed for translation machinery components (i.e. RplU [Tyr4], and RpsE [Ser159]), Pgi (Thr149), SMU_1080c (Thr47), SMU_502 (Thr52 and Ser53), and SMU_668c (Ser157). We suppose that these changes are PknB-independent.

**Figure 7:**
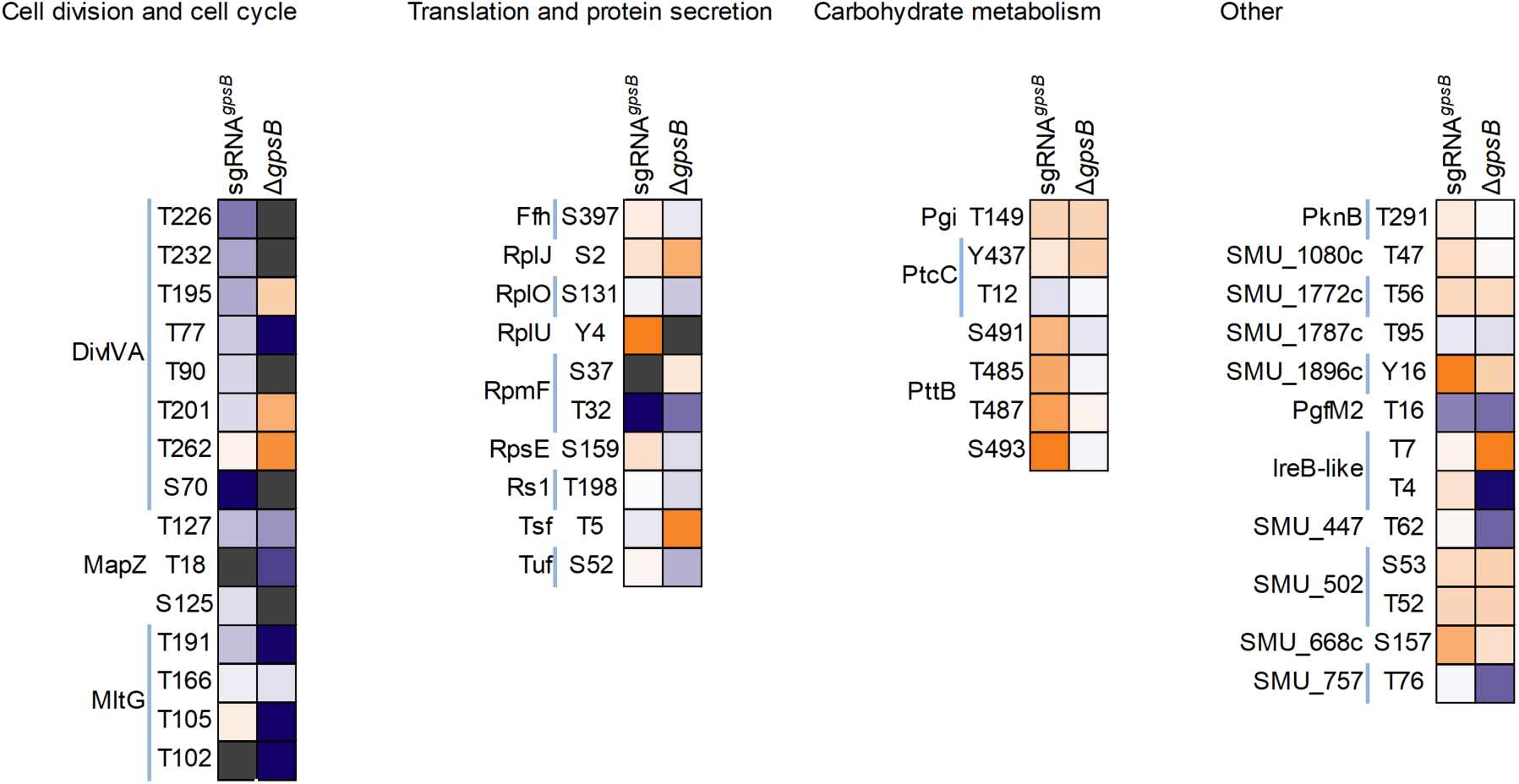
Heatmap visualizing hyper- and hypophosphorylation of serine, threonine, and tyrosine residues in CRISPRi*^gpsB^* and Δ*gpsB*_sup_ *S. mutans* strains. Phosphorylated proteins are sorted according to their biological function. Orange indicates increased phosphorylation compared to the control strain, and blue indicates reduced phosphorylation compared to the control strain. Grey indicates that phosphorylation was not quantified at that residue for that strain.

As described, the CRISPRi*^gpsB^* strain has severe phenotypes that are apparently restored in the Δ*gpsB*_sup_ strain. We posit that restoration of growth is caused by a reversal in the phosphorylation phenotype that we have observed in the CRISPRi*^gpsB^* strain. PhosphoTMT analysis of Δ*gpsB*_sup_ revealed clear differences compared to the CRISPRi*^gpsB^* strain. Notably, there was a significant increase in phosphorylation (p-value < 0.05; log2 fold-change > 1) at Thr195, Thr201, and Thr262 in DivIVA. This increase was residue-specific for DivIVA as phosphorylation was absent at Thr77 and was also significantly decreased (p = 0.001; log2 fold-change = −1.32) in the CRISPRi*^gpsB^* strain. Increased phosphorylation of DivIVA was the most notable restoration of phosphorylation among the putative PknB cellular substrates. MapZ (Thr127, −2.6), MltG (Thr105, absent; Thr191, −5.9), RpmF (Thr32, −3.4), PgfM2 (Thr16, −3.5), and SMU_2079c (Thr4, −5.8; Thr7, 4.8) still displayed significantly decreased phosphorylation in the CRISPRi*^gpsB^* strain. This raises the possibility that *gpsB* essentiality is caused by aberrant phosphorylation of DivIVA. In *S. mutans* UA159, *divIVA* is not an essential gene, as determined by Tn-seq (32). Our results could suggest that altered phosphorylation of DivIVA does not remove the function of the protein but instead causes it to behave in a way that is deleterious to the cell. Other notable alterations in phosphorylation, compared to CRISPRi*^gpsB^*, were observed for Tsf (Thr5, 4.6), SMU_447 (Thr62, −3.7), and SMU_757 (Thr76, −3.8).

## Discussion

In this study, we provide the first comprehensive analysis of *O*-phosphorylation in *S. mutans*. It highlights the significant roles of both the kinase PknB and the phosphatase PppL in regulating key biological processes like translation, carbohydrate metabolism, and the cell cycle. Of the putative PknB and PppL targets, some are unique to *S. mutans,* and others overlap with targets in other Gram-positive microorganisms. Our study also identified a novel suppressor mutation in *pppL* that can bypass the essentiality of *gpsB* in *Streptococcus mutans*, a phenomenon previously observed in other Gram-positive bacteria with different phosphatase mutations. While prior research focused on the role of GpsB in protein phosphorylation, this work uniquely demonstrates that the G98R mutation in PppL leads to hyperphosphorylation, specifically restoring phosphorylation of DivIVA, which appears crucial for reversing the severe phenotypes associated with *gpsB* repression. Taken together, our data suggests a nuanced interplay between GpsB, PknB, and PppL in modulating cell division and other critical pathways in *S. mutans*.

Although bacterial phosphoproteomes are generally less complex than those of eukaryotes, our data demonstrate that *S. mutans* exhibits a substantial degree of protein phosphorylation that is consistent with findings in *S. pneumoniae* (4% of total proteins) (33), *S. thermophilus* (5.8%) (34), *B. subtilis* (3%) (35), and *L. monocytogenes* (6.6%) (36). In addition, the relative abundance of phosphothreonine, phosphoserine, and phosphotyrosine is consistent with prior observations in other Gram-positive species. Despite the overall less frequent detection of phosphotyrosine, this observation is interesting as the *S. mutans* genome lacks a BYK. This raises the possibility that a novel tyrosine kinase remains to be identified, or that PknB has dual-specificity that includes tyrosine. Dual-specificity protein kinases, that phosphorylate serine, threonine, or tyrosine, have been detected in bacterial species but these are distinct from STKPs (37). Our own genome mining analysis found that SMU_65 (WP_002263403.1) encodes for a low molecular weight protein tyrosine phosphatase (LMWPTP). This could be a key regulator of tyrosine phosphorylation that warrants further investigation, along with the identification of a protein with tyrosine kinase activity.

The TMT-labeling, IMAC enrichment, and MaxQuant analysis pipeline represents the gold standard for phosphoproteomic profiling. However, the biological significance of the observed phosphorylation events—particularly those occurring on essential proteins such as MapZ, GpsB, and ribosomal components—remains to be elucidated through functional follow-up studies. Of particular interest are six proteins whose phosphorylation appears to be PknB-dependent: DivIVA, MapZ, RpmF, MltG, an IreB-like protein (SMU_2079c), and PgfM2. Among these, DivIVA, MapZ, MltG, and the IreB-like protein have been previously identified as substrates of PknB, while RpmF and PgfM2 represent novel candidates. PgfM2 is a compelling target for future investigation. It encodes a highly conserved core gene in *S. mutans* and functions as part of the Pgf glycosylation machinery (20). In the *S. mutans* strain OMZ175, the Pgf system is responsible for glycosylating several critical proteins, including the collagen-binding adhesin Cnm (20, 38, 39). Despite its established importance in OMZ175, the role of PgfM2 in the widely studied laboratory strain UA159 remains less well understood. Our phosphoproteomic data suggest potential cross-talk between protein phosphorylation and glycosylation systems. This raises the intriguing possibility that phosphorylation of PgfM2 may regulate its function or the broader activity of the Pgf machinery. Functional experiments will be necessary to determine the biological consequences of PgfM2 phosphorylation and to clarify the mechanistic basis of the linkage between these two post-translational modification systems. In Gram-positive bacteria, glycosylation modifies multiple cell wall-associated polymers, including rhamnose-rich polysaccharides and lipoteichoic acids (40). The PASTA domain of STKP proteins is known to respond to extracellular peptidoglycan muropeptide fragments (41). Thus, the phosphorylation of PgfM2 by PknB may represent a regulatory intersection at the cell envelope, potentially integrating signals that coordinate cell wall remodeling with cell wall polymer glycosylation.

Our findings support a model in which GpsB is a modulator of the PknB/PppL signaling axis, ensuring appropriate phosphorylation of key cell division and cell wall biosynthesis proteins. Loss of GpsB leads to hypophosphorylation of essential substrates and cell lethality, but this can be genetically compensated by mutations that impair PppL activity and restore phosphorylation levels. This interplay reveals a finely tuned regulatory network where phosphorylation states of a select group of proteins are critical for cell viability, and where GpsB and PppL act as central nodes. Our findings in *S. mutans* parallel those in other Gram-positive microorganisms with some key species-specific differences. A key similarity is the observation that *S. pneumoniae phpP* mutations that inactivate phosphatase activity allow *gpsB* deletion (30). While the mutations recovered in *phpP* are different to the G98R mutation we recovered, the broad observation is the same. In *S. mutans*, five of six putative PknB cellular substrates— DivIVA, MapZ, MltG, RpmF, and PgfM2—showed reduced phosphorylation upon *gpsB* repression, indicating that GpsB is required for efficient PknB-mediated phosphorylation of these targets. This is consistent with prior findings in *S. pneumoniae*, where GpsB appears to support PknB activity and/or substrate localization (24). Interestingly, the phosphorylation profile of PknB itself remained unchanged in both the *gpsB*-repressed and deleted strains, suggesting that GpsB is not required for PknB autophosphorylation, in contrast to proposed models in other species (26). Compared to other studies there is divergence of PknB and GpsB interplay regarding PknB-mediated phosphorylation of GpsB. We were unable to observe a decrease in GpsB phosphorylation in a *pknB* mutant background whereas STKPs have been shown to be required for GpsB phosphorylation both in *B. subtilis* and *S. pneumoniae* (22, 23). However, evidence of this interaction is mixed in *S. pneumoniae* as StkP did not appear to phosphorylate GpsB in a different strain background (D39 vs. Rx1) under different conditions (42). This interaction could be complex, and we cannot rule out phosphorylation of GpsB by PknB being influenced by growth phase or environmental conditions. In *Clostridioides difficile*, for example, phosphoproteome profiles change dramatically depending on growth phase (43). It is therefore plausible that GpsB phosphorylation by PknB in *S. mutans* might occur under specific stress or developmental signals not captured under our experimental conditions.

Our data suggests that the essentiality of *gpsB* in *S. mutans* may arise from a failure to maintain proper phosphorylation of DivIVA, resulting in deleterious alterations to its function. While GpsB may not be inherently essential, its absence appears to cause lethal dysregulation of DivIVA activity through altered phosphorylation. Among the putative PknB substrates analyzed, DivIVA showed the most pronounced recovery of phosphorylation in the Δ*gpsB*_sup_ strain, whereas others such as MapZ, MltG, RpmF, and PgfM2 remained hypophosphorylated. This highlights a potentially unique and central role for DivIVA in mediating the suppressor phenotype. DivIVA serves diverse functions across the Bacillota, including roles in cell division, chromosome segregation, and cell wall synthesis, with both conserved and species-specific activities (reviewed in Hammond, White, and Eswara (21)). In *S. pneumoniae*, DivIVA coordinates septum positioning and peripheral peptidoglycan synthesis (44), interacting with key proteins such as FtsZ and MltG. It is plausible that disrupted phosphorylation of DivIVA impairs these protein-protein interactions. Supporting this, bacterial two-hybrid studies in *Streptococcus suis* have shown that phospho-defective and phospho-mimetic variants of DivIVA exhibit altered interactions with MltG (45). Moreover, phosphorylation may regulate DivIVA turnover, as previous work indicates it can be targeted for ClpXP-mediated degradation (46). Taken together, these findings support a model in which aberrant DivIVA phosphorylation converts it from a functional scaffold to a toxic entity, potentially through misregulation of cell division and peptidoglycan synthesis. Future studies employing phospho-null and phospho-mimetic DivIVA mutants will be critical for determining whether phosphorylation state alone is sufficient to drive viability or lethality in *gpsB*-deficient backgrounds.

Taken together, our findings reveal a finely tuned phosphorylation network governing *S. mutans* physiology, with species-specific features that parallel yet diverge from established paradigms in other Gram-positive bacteria. This work lays the groundwork for future functional studies aimed at dissecting how specific phosphorylation events impact cellular function, and how cross-talk between post-translational modifications integrates environmental signals into the regulation of bacterial growth, division, and pathogenic potential.

## Materials and Methods

### Bacterial strains and culture conditions

*Streptococcus mutans* strains were cultured from single colonies in Brain Heart Infusion (BHI, Difco) broth. Unless otherwise noted, *S. mutans* was routinely cultured at 37°C in a 5% CO_2_ microaerophilic atmosphere. Antibiotics were added to growth media at the following concentrations: kanamycin 1 mg/mL, and spectinomycin 1 mg/mL. A list of strains (**Table S11**) can be found in the supplementary material.

#### Construction of strains

A PCR ligation mutagenesis method was used to replace genes and non-coding regions with non-polar kanamycin markers (47). For each gene deletion, primers A (GGAGAGCACTGGAGTGAAGC) and B (ATGCGGATCCCATAGCCACGCATGCTAGATT) were designed to amplify 500-600 bp upstream of the coding sequence (with ca. 50-bp overlapping the coding sequence of the gene). Primers C (ATGCGGATCCTGGCGACTACCGCTACAAAT) and D (TTGTTCCAAAGGCGACTAGG) were designed to amplify 500 to 600 bp downstream of the coding sequence (with ca. 50-bp overlapping the coding sequence of the gene). Primers B and C contained BamHI restriction enzyme sites for ligation of the AB and CD fragments to a non-polar kanamycin cassette digested from plasmid pALH124 (30). Transformants were selected on BHI agar containing kanamycin. Double-crossover recombination, without introduction of nearby secondary mutations, was confirmed by PCR and genome sequencing.

### Genome sequencing

Genomic DNA was isolated from strains using a MasterPure Gram Positive DNA (Epicentre) purification kit with modifications as previously described (48). After DNA purification, total DNA concentration and purity was measured using a Qubit dsDNA Broad Range Assay Kit (Thermo Fisher Scientific). DNA samples (total volume of 30 µL; > 10 ng/ µL) were sent to SeqCenter (Pittsburgh, USA) for whole genome sequencing. We ordered the 200 Mbp service, with variant calling compared to the *S. mutans* UA159 genome (NC_004350.2).

### Tandem mass tag mass spectrometry

*S. mutans* cells were grown to mid-log phase (∼0.4-0.6 OD_600_) in 15 mL of growth medium, and cell pellets were subsequently harvested. Five biological replicates were collected for each strain. The cell pellets were shipped on dry ice to the IDeA National Resource for Quantitative Proteomics at the University of Arkansas for Medical Sciences (UAMS; Little Rock, USA). Protein extraction was performed using chloroform/methanol, followed by trypsin digestion. After digestion, individual samples were labeled with TMT tags. Phosphorylated proteins were enriched using immobilized metal ion affinity chromatography (IMAC) resins. Due to the high complexity of these samples, the peptide mixtures underwent reverse-phase chromatography fractionation to simplify them and enhance phosphoproteome coverage. Samples were then loaded onto an autosampler, concentrated with a trapping column, and separated by an acetonitrile gradient on an analytical column. The separated peptides were analyzed by MS and MS/MS using an Orbitrap Eclipse (Thermo Scientific) platform. MaxQuant, a quantitative proteomics software, was used to search for *S. mutans* UA159 proteins. Quality control, normalization, and differential expression analysis were conducted afterward.

### Transmission electron microscopy

Bacterial cultures were grown overnight to stationary phase with appropriate concentrations of xylose (w/v) in a chemically defined medium (CDM) (49) to induce CRISPRi repression. The culture samples were centrifuged at 3,500 x *g* for 5 min to produce a cell pellet, which was washed twice using phosphate buffered saline for 10 min, with 2 min of centrifugation at 12,000 x *g* between each wash. Following removal of the media and PBS buffer, 1 mL of glutaraldehyde was added to fix the sample. Samples remained in fixative at least overnight, then fixative was removed and replaced with 1 mL of PBS before being sent to the UAMS Digital Microscopy Core (Little Rock, USA) for TEM processing. At UAMS, cells were treated with 1.5% osmium tetroxide in the dark for 1 h at 4°C. Then the cells were mixed in equal parts with 5% agarose in PBS, collected by centrifugation at 2000 x g, and cooled to 4°C. Small chunks of the bacterial pellet plus agarose were then dehydrated in ethanol via the following steps: 30%, 50%, 70%, 80%, 90% each for 15 min, 99% 10 min, and then absolute ethanol 2 x 10 min. The dehydrated cells were embedded in an epoxy resin, sectioned and stained with uranyl acetate and lead citrate. Microscopy was conducted using a FEI Tecnai F20 transmission electron microscope.

### RNA isolation and RNA sequencing

Bacterial strains were cultured in CDM to an OD_600_ ∼0.4–0.6 and treated with RNAprotect (Qiagen). Cell pellets were resuspended in a lysis buffer (50 mM Tris-HCl, pH 8.0, 10 mM EDTA, and 0.4% SDS). Next, the cell suspensions were placed in screw-cap 2 mL tubes containing 100 mg of 0.1 mm glass beads and mixed with an equal volume of acidic phenol:chloroform:isoamyl alcohol (pH 5.2; 25:24:1; 1:1, vol/vol). After bead beating using a Mini-Beadbeater-24 (BioSpec), the sample tubes were centrifuged for 5 mins at 3500 x *g* and the clarified aqueous layer was removed. RNA was then extracted using the RNeasy Mini Kit (Qiagen), with genomic DNA removal in column using RNase-free DNase I solution (Qiagen). RNA was quantified using a Qubit RNA Broad Range Assay Kit (Thermo Fisher). Next, RNA was sent to SeqCenter who generated RNA-seq libraries using Illumina stranded RNA library preparation with RiboZero Plus rRNA depletion. Sequencing was conducted on an Illumina platform providing up to 12 million paired end reads (2×51 bp) per sample. After sequencing, SeqCenter conducted a basic RNA-seq analysis pipeline that provided raw transcript level quantification. Afterwards, gene expression changes between samples were quantified with Degust (http://degust.erc.monash.edu/) using the edgeR methodology.

## Acknowledgements

This work was supported by the Arkansas INBRE program, supported by a grant from the National Institute of General Medical Sciences (NIGMS), Grant P20 GM103429 (R.C.S.) from the National Institutes of Health.

